# Adult-born neurons promote cognitive flexibility by improving memory precision and indexing

**DOI:** 10.1101/2020.08.10.242966

**Authors:** Gabriel Berdugo-Vega, Chi-Chieh Lee, Alexander Garthe, Gerd Kempermann, Federico Calegari

## Abstract

The dentate gyrus (DG) of the hippocampus is fundamental for cognitive flexibility and has the extraordinary ability to generate new neurons throughout life. Recent evidence suggested that adult-born neurons differentially modulate input to the DG during the processing of spatial information and novelty. However, how this differential regulation by neurogenesis is translated into different aspects contributing cognitive flexibility is unclear. Here, we increased adult-born neurons by a genetic expansion of neural stem cells and studied their influence during navigational learning. We found that increased neurogenesis improved memory precision, indexing and retention and that each of these gains was associated with a differential activation of specific DG compartments and better separation of memory representations in the DG-CA3 network. Our results highlight the role of adult-born neurons in promoting memory precision in the infrapyramidal and indexing in the suprapyramidal blade of the DG and together contributing to cognitive flexibility.

**One sentence summary:** Neurogenesis improves memory precision and indexing.

## INTRODUCTION

The hippocampal DG-CA3 network is fundamental for the separation and recollection of episodic memories (Rolls, 2013). In addition, this circuit possesses the exceptional ability to incorporate new DG granule cells over the course of life through a process called adult neurogenesis. Adult neurogenesis has been proposed to promote cognitive flexibility, which is defined as the ability to adapt learning to the presence of changing contingencies (Anacker and Hen, 2017; Garthe et al., 2009). This notion was supported by a number of loss of function studies (Burghardt et al., 2012; Clelland et al., 2009; Dupret et al., 2008; Garthe et al., 2009; Niibori et al., 2012) as well as approaches increasing neurogenesis and resulting in improvements in hippocampal-dependent learning in reversal paradigms of spatial navigation and contextual discrimination (Berdugo-Vega et al., 2020; Epp et al., 2016; Garthe et al., 2016; Kent et al., 2015; McAvoy et al., 2015; Sahay et al., 2011; Wang et al., 2014). Importantly, several complex cognitive processes can independently improve flexible learning and contextual discrimination. Among these, mathematical models proposed a role of adult-born neurons in resolving memory interference (Appleby et al., 2011; Wiskott et al., 2006). In parallel, experimental manipulations suggested that such neurogenesis-dependent resolution of memory interference can be achieved either through an increased sparsity of neural representations (McAvoy et al., 2015), their consolidation (Kitamura et al., 2009) or, alternatively, forgetting (Akers et al., 2014). These different aspects of cognitive flexibility were intrinsically difficult to disentangle and the specific role(s) of adult-born neurons in memory precision, separation or clearance remained elusive.

Furthermore, adult-born neurons were proposed to either act as encoding units or regulators of the activity of the more numerous, mature granule cells (Piatti et al., 2013). Independently from their mode of action, effects of neurogenesis were generally assumed to be homogeneous within the DG. However, recent studies highlighted that adult-born neurons differentially modulate input from the entorhinal cortex into the DG (Luna et al., 2019), which is further supported by an intrinsic heterogeneity in the cellular composition of its infra-vs. supra-pyramidal (IP and SP, respectively) blades (Erwin et al., 2020; Mishra and Narayanan, 2020). Intriguingly, the IP and SP blades are known to preferentially receive input about spatial information or novelty that originate from the medial or lateral entorhinal cortex, respectively (Hargreaves et al., 2005; Knierim et al., 2014; Luna et al., 2019). This anatomical distinction made us speculate that a differential modulatory effect of adult-born neurons within the two blades of the DG may support different aspects of cognitive flexibility. In turn, such anatomical and functional separation may help reconcile conflicting studies proposing a role of neurogenesis in memory precision, separation or, alternatively, forgetting.

Here, we sought to dissect the different components of neurogenesis-dependent, cognitive flexibility and their anatomical allocation. To this end, we exploited a mouse model in which increased neurogenesis was achieved by a Cdk4/cyclinD1-dependent expansion of neural stem and progenitor cells (together NSC) (Artegiani et al., 2011) that was shown to rescue hippocampal cognitive deficits during aging (Berdugo-Vega et al., 2020). This approach gave us the unique opportunity to investigate the impact of neurogenesis upon generation of an expanded and age-matched cohort of adult-born neurons, thus, avoiding confounding effects resulting from chronic treatments, such as physical exercise or irreversible genetic manipulation, in which neurons of different ages can simultaneously trigger different behavioral effects. We further combined this genetic manipulation of neurogenesis with engram labelling technologies used as a proxy for the identification of behaviourally-induced neuronal activity (Tonegawa et al., 2015) and assessed memory ensembles in the IP vs. SP blade during the processing of spatial information vs. novelty. Furthermore, memory representations and their overlap were identified to investigate the role of adult-born neurons in memory precision, separation or clearance and their anatomical allocation. Together, our study aimed to dissect different aspects of adult neurogenesis in the refinement of memory indexes and cognitive flexibility.

## RESULTS

### Transient expansion of NSC results in the generation of a single, age-matched cohort of newborn neurons in the IP and SP blades

Our group already described a method to increase neurogenesis by a genetic expansion of NSC. This was based on the overexpression of Cdk4/cyclinD1 (4D for brevity) in NSC by means of stereotaxic injection of lentiviral particles in the DG. Achieving the temporal control of NSC expansion, a loxp-flanked 4D cassette was used and injections performed in *nestin*::CreERt2 mice. In essence, while viral injection led to overexpression of 4D and NSC expansion, tamoxifen was administered 3 weeks later to terminate this effect and induced the synchronous switch of the expanded cohort of NSC to increased neurogenesis (Artegiani et al., 2011). Having shown the efficacy of this approach (Berdugo-Vega et al., 2020), and similar strategies resulting from it (Bragado Alonso et al., 2019), we here sought to exploit this tool to generate a single wave of age-matched newborn neurons of 4 weeks of age, a timepoint when they are known to display functional integration and enhanced synaptic plasticity before becoming undistinguishable from mature granule cells (Esposito et al., 2005; Gu et al., 2012; Restivo et al., 2015; Temprana et al., 2015). Hence, neuronal maturation, survival and integration were assessed in 4D-injected mice (2-3 months old) at different times after tamoxifen administration with specific focus on their distribution in the IP and SP blades of the DG.

First, we took advantage of the design of our viral vectors triggering redistribution of a GFP reporter from the nucleus to the cytoplasm upon Cre recombination in NSC (in both control and 4D) and assessed the morphology of newborn neurons at different times (2, 4 and 6 weeks) after tamoxifen administration (Fig. 1A). This showed a comparable progression in the complexity of dendritic arborization, spine density and axonal projections to the CA3 in GFP-control and 4D-derived newborn neurons over time (Fig. 1B, S1A). Together with previous reports from our group (Berdugo-Vega et al., 2020; Bragado Alonso et al., 2019), these results validated the physiological differentiation and integration of newborn neurons upon 4D overexpression in NSC. At the same time, these experiments also gave us the opportunity to assess whether the increase in neurogenesis triggered by the expansion of NSC was constant or transient. Comparison of the levels of putative NSC (Sox2+S100β–) and early born neurons identified by a marker transiently expressed after birth (Dcx+) within the pool of virally transduced cells (GFP+; Fig. S1B) confirmed a 2-fold increased levels of neurogenesis in 4D-injected mice 2 weeks after tamoxifen (Sox2+S100β–: 8.51 ± 1.56 vs. 3.95 ± 1.67%, *p* = 0.026; Dcx+: 6.76 ± 1.16 vs. 3.57 ± 0.96%, *p* = 0.021 for 4D and GFP mice, respectively; Fig. 1C, left). In contrast, both NSC and early born neurons returned to physiological levels 4 weeks post tamoxifen (Sox2+S100β–: 4.80 ± 1.90 vs. 4.27 ± 0.25%, *p* = 0.66; Dcx+: 3.69 ± 0.67 vs. 4.15 ± 2.53% for 4D and GFP, respectively, *p* = 0.77; Fig. 1C, right) indicating that the increase in neurogenesis triggered by 4D was transient. Yet, the abundance of total 4D-derived neurons identified by cytoplasmic GFP remained the double at 2 and 4 weeks after tamoxifen (4D relative to GFP, fold increase: 2.71 ± 0.38 and 2.26 ± 0.66, respectively, *p* = 0.012 and 0.033; Fig. 1C) suggesting that the expanded wave of age-matched newborn neurons was not compensated by increased cell death. This was confirmed by BrdU birthdating one week prior to tamoxifen administration and assessment of adult-born neurons 4 weeks later by the use of a mature marker (NeuN+). This showed that the wave of an increased cohort of neurons was maintained in 4D animals with a 2-fold increase in 4 week old neurons relative to controls (NeuN+BrdU+: 0.64 ± 0.09 vs. 0.25 ± 0.05%, *p* = 0.003) that was of comparable magnitude in the IP and SP blades (2.12-fold, *p* = 0.044 and 3.28-fold, *p* = 0.013, respectively) and similar among the two groups of mice (IP vs. SP within GFP and 4D: *p* = 0.53 and *p* = 0.65, respectively; Fig. 1D).

**Figure 1:**
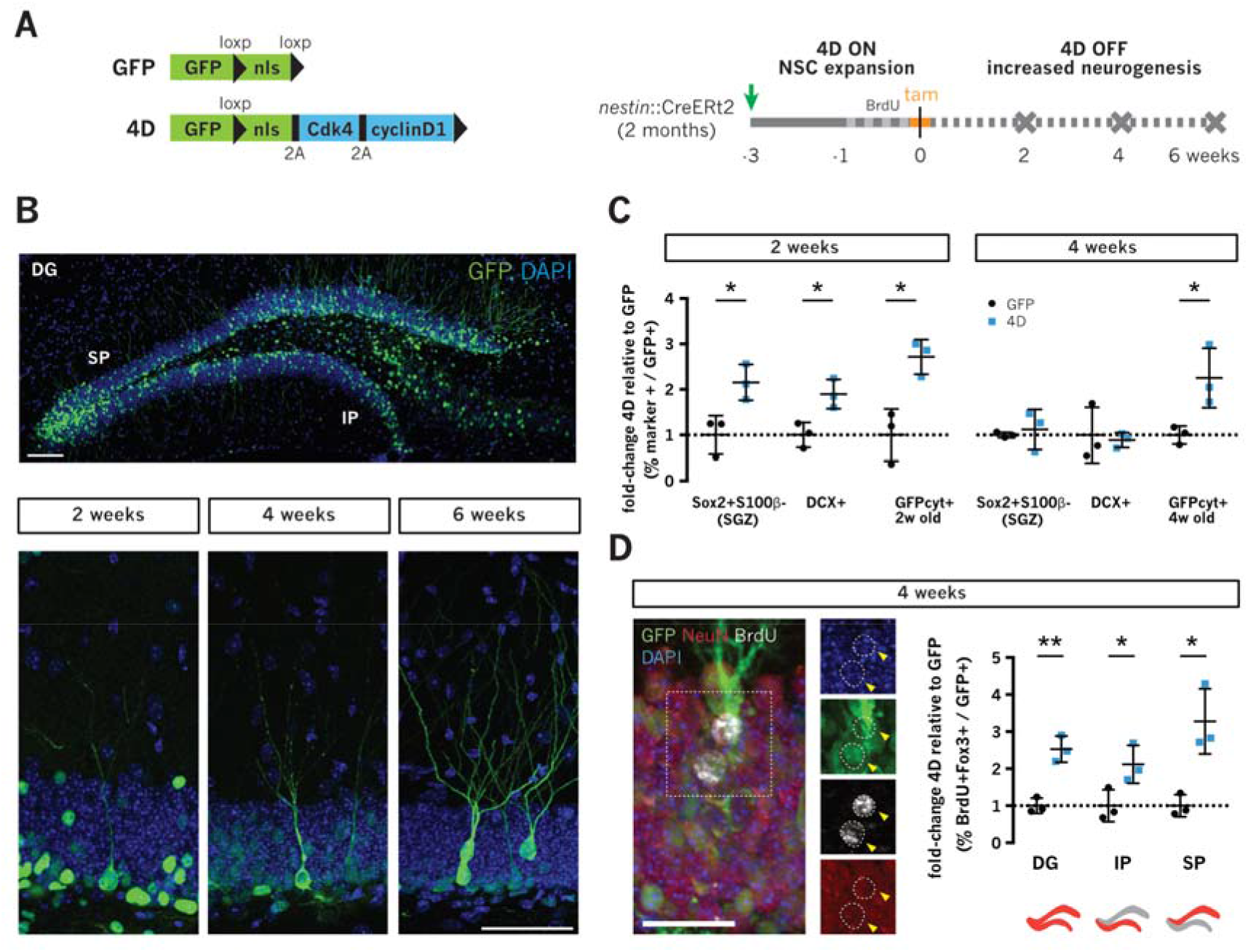
NSC expansion generates a single, age-matched cohort of newborn neurons in the IP and SP blades. **A)** Scheme depicting the GFP and 4D lentiviral constructs (left) and experimental layout (right) to achieve temporal control of transgenes expression by injection of lentiviruses in the DG of 2-3 month old *nestin*::CreERt2 mice (4D ON) with tamoxifen triggering the recombination of the loxp-flanked functional genes (4D OFF) together with the nuclear localization signal (nls) of GFP. **B-D**) Fluorescence pictures and quantifications of cell types identified by immunohistochemistry in the subgranular zone (SGZ) of the DG (C) and IP or SP blades (D) at different times after tamoxifen administration (as indicated). Note the nuclear to cytoplasmic redistribution of GFP in newborn neurons (B; bottom). Examples of cells scored are indicated (D, insets; arrowheads). Scale bars = 100 μm (B) and 50 μm (D). Data represent mean ± SD as fold-change within GFP+ infected cells of 4D (blue) relative to GFP (black). N = 3; n > 500 (SGZ) and > 1,000 (IP/SP). Unpaired Student’s t-test \**p* < 0.05; ** *p* < 0.01.

Together, our data showed that a transient expansion of NSC can be used to generate a synchronized neurogenic wave in the IP and SP. Several loss of function studies highlighted the effects of ablating neurogenesis in negatively influencing hippocampal function (Burghardt et al., 2012; Clelland et al., 2009; Dupret et al., 2008; Garthe et al., 2009; Niibori et al., 2012). Our approach gave us the unique opportunity to assess the impact of the converse manipulation upon generation of an expanded and age-matched cohort of 4 week old newborn neurons.

### Increased neurogenesis promotes spatial navigation and flexible learning by differentially modulating the activity of the IP and SP

We next investigated the effect of this increased wave of 4 week old neurons in mouse cognitive function by using a paradigm of the Morris water maze including a learning phase followed by reversal to assess spatial navigation and novelty processing, respectively (Fig. 2A). Our group recently showed that, in aged mice, assessment of navigational learning strategies revealed important aspects of cognition that would not necessarily be apparent when assessing behavioral performance alone (Berdugo-Vega et al., 2020). Hence, we tested GFP and 4D-injected mice 4 weeks after tamoxifen administration to evaluate their evolution in the use of hippocampus-dependent and spatially-precise strategies over time by using an automated machine learning model that assesses swimming trajectories (Overall et al., 2020).

**Figure 2:**
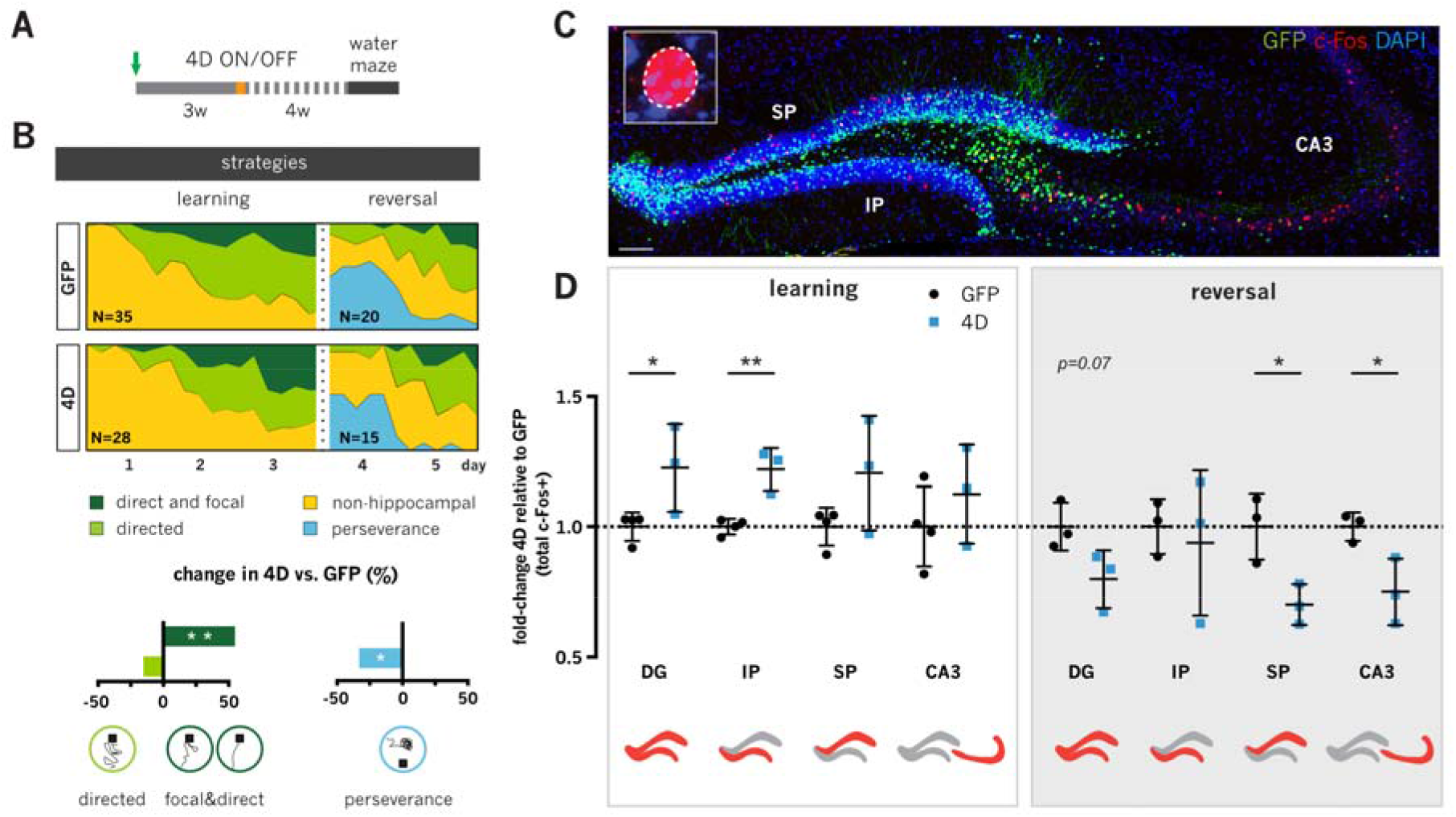
Increased neurogenesis promotes spatially precise navigation and flexible learning by differentially modulating the activity of the IP and SP blades. **A-D)** Experimental layout (A) and evolution in navigational strategies (B) of GFP and 4D-treated mice subjected to a water maze test after treatment shown in A) and sacrificed after the last trial of learning (day 3) or reversal (day 5) for assessment the of c-Fos (C; magnified in inset) in the DG/IP/SP/CA3 compartments (D; as indicated). The same data are also depicted as a comparison between learning vs. reversal within groups of GFP or 4D mice in Fig. S2C. Note the increased use of direct and focal strategies in the last learning day with reduced perseverance after reversal (B; dark green and blue, respectively) correlating with increased c-Fos in the IP blade (D; left) and decrease in the SP-CA3 (D; right) in 4D (blue) relative to control (black) mice. Scale bar = 100 μm. Data represent mean ± SD as fold-change of c-Fos+ cells in different DG-CA3 compartments. N = 15-35 (B) or 3-4 (D) mice; n > 50 (IP/SP) and > 130 (CA3). Wald test (B) or unpaired Student’s t-test (D) * *p*<0.05; ** *p*<0.01.

During the learning phase, the increase in the use of hippocampus-dependent strategies at the expense of non-spatial navigation progressed at a similar pace and to a similar degree in both control and mice with increased neurogenesis (Fig. 2B). This was in contrast to the recently described effect of increasing neurogenesis in old mice, in which 4D rescued age-dependent deficits by promoting hippocampal navigation (Berdugo-Vega et al., 2020). However, since young mice likely already display the highest levels of hippocampal navigation, we investigated whether the precision of these swimming strategies were different on the last day of the learning phase. Mice with increased neurogenesis showed more direct trajectories (initial trajectory error: 48.7 ± 16.7 vs. 58.7 ± 16.2 cm for 4D and GFP respectively, *p* = 0.020) and shorter pathlengths among hippocampally scored strategies (311.0 ± 167.2 vs. 357.7 ± 165.6 cm for 4D and GFP respectively, *p* = 0.002; Fig. S2B). We then used the pathlength as a proxy of precision and divided hippocampal strategies in spatially precise or imprecise (pathlengths <1.5 or >1.5 over optimal performance, respectively) and finding in mice with increased neurogenesis a 2.3-fold increase in precise strategies relative to controls (logistic regression, odds ratio [OR] = 3.02, *p* = 0.0001; Fig. S2B). Confirming this, the previous unbiased assessment of swimming trajectories revealed a 58% increase in the proportion of “direct” and “focal” strategies in 4D relative to control mice (together: OR = 2.19, *p* = 0.002; Fig. 2B, bottom). Hence, these results indicated that while increasing neurogenesis did not seem to influence the pace of the switch to hippocampal learning strategies, it nonetheless improved their precision once these were acquired.

We then investigated neuronal correlates of spatial memory by performing c-Fos immunohistochemistry of the dorsal hippocampus (Fig. 2C). Analysis of mice killed after the last learning trial revealed an increased activity of the DG of 4D-treated mice relative to controls (total estimated c-Fos+ per hemisphere: 8,034 ± 1,102 vs. 6,556 ± 359, respectively, *p* = 0.049; Fig. 2D, left). Specifically, this was due to a selective increase by 20% in the number of active granule cells in the IP blade (2,496 ± 167 vs. 2,046 ± 62 for 4D and GFP, respectively, *p* = 0.004) while values in the SP were not significantly changed (Fig. 2D, left). This was consistent with the role of newborn neurons in increasing neuronal activity during spatial navigation (Luna et al., 2019). It also suggested that the recruitment of a larger neuronal ensemble specifically within the IP blade improved navigation and memory precision.

To further address the effect of increased neurogenesis upon the introduction of novelty, a group of mice was additionally trained for two days upon reversal of the platform’s position to the opposite quadrant of the pool. Analysis of the swimming trajectories showed the appearance of perseverance in both groups and reacquisition of hippocampal-directed strategies thereafter (Fig. 2B). Specifically, the contribution of perseverance (blue) to navigational strategies was significantly reduced by 30% in mice with increased neurogenesis (OR = 0.62, *p* = 0.032; Fig. 2B; bottom), which is indicative of improved flexible learning (Garthe et al., 2009). This showed that the more spatially precise, 4D mice were able to more flexibly adapt after the introduction of novelty.

We next performed a similar assessment of c-Fos in mice after the last trial of reversal. Here, 4D-treated mice showed a general decrease in the activity of the DG (c-Fos+: 6,293 ± 876 vs. 7,884 ± 727, for 4D and GFP, respectively, *p* = 0.072) that was specifically due to a 30% reduction in the number of activated cells in the SP blade (3,679 ± 406 vs. 5,255 ± 671 in 4D and GFP respectively, *p* = 0.025) without significant changes in the IP blade (Fig. 2D, right). In addition, quantification after reversal showed the first changes in activity within the CA3 with a decrease by 25% relative to controls (5,996 ± 1,015 vs. 8,002 ± 439, *p* = 0.035; Fig. 2D, right) that was instead not detected in the previous quantifications of the learning phase (7,031 ± 1,189 vs. 6,251 ± 959 for 4D and GFP respectively, *p* = 0.37; Fig. 2D, left).

These experiments also gave us the opportunity to reanalyse the same data, but this time considering the intra-group analysis within GFP or 4D mice after learning and reversal. Within the two groups of GFP mice, comparison of c-Fos before and after reversal revealed a general increase in the overall activity of the DG due to elevated counts in the IP blade (*p* = 0.006) and accompanied by a similar trend in the SP (*p* = 0.13; Fig. S2C, left). Notably, in control mice a comparable increase in c-Fos was also found in CA3 (*p* = 0.034; Fig. S2C, left). Conversely, intra-group comparisons of 4D mice showed that their response to novelty resulted in a general reduction in DG activity through a specific decrease of the SP blade (*p* = 0.043; Fig. S2C, right). Additionally, no correlation in the activity of the DG and CA3 was detected after learning (r = −0.03, *p* = 0.95) while a strong positive correlation appeared in both groups after reversal (r = 0.89, *p* < 0.0001; Fig. S2D) suggesting the active involvement of this circuit in novelty detection.

Consistent with a differential modulation of entorhinal input by newborn neurons (Luna et al., 2019), our results show that increased neurogenesis elevated the activity specifically of the IP and that this was associated with the use of more spatially precise strategies during learning indicative of higher memory precision. Additionally, we found that changes in flexible learning were associated with a differential response to the introduction of novelty, with control mice increasing DG-CA3 activity while 4D mice decreased it. Notably, none of these changes directly involved the newborn neurons themselves since these where never found to be part of any c-Fos+ ensemble (data not shown). Knowing that neurogenesis-mediated sparsity was suggested to reduce memory interference (Anacker and Hen, 2017; McAvoy et al., 2015), we next investigated whether animals with increased neurogenesis created new and more separated memory representations in the SP-CA3 circuit after reversal.

### Cognitive flexibility without forgetting is associated with a better separation of memory indexes in the SP-CA3 circuit

To directly assess whether neurogenesis-dependent downregulation of SP activity alone decreased the overlap of memory representations, we sought to identify the ensemble of neurons representing the first learning experience and compare it with that formed after reversal by the use of engram-labelling technology (Reijmers et al., 2007).

GFP and 4D-treated mice were maintained in doxycycline (dox) diet and additionally injected prior to tamoxifen administration with AAV9 viruses encoding *c-Fos*::tTa and *TRE*::mCherry (Fig. 3A). Mice were trained in the water maze and dox removed from the diet 24 h before the last day of the learning phase to allow the activity-dependent labelling of the memory ensemble representing the first experience. Immediately after the last trial of learning, dox was reintroduced and mice subjected to two additional days of reversal and killed after the last trial (Fig. 3A). The reactivation rate was finally assessed as defined by the proportion of cells belonging to the remote memory ensemble (mCherry+) that were also recruited by the recent memory (c-Fos+ revealed by immunohistochemistry; Fig. 3B and S3A). This showed that reactivation in the DG of 4D mice was ca. 30% lower than in controls (2.25 ± 0.42 vs. 3.30 ± 0.56%, respectively, *p* = 0.043), which was selectively due to a ca. 40% reduction in the SP blade (2.17 ± 0.40 vs. 3.66 ± 0.53, *p* = 0.010; Fig. 3C). Interestingly, a similar decrease by 44% in reactivation rate was also observed in the CA3 of 4D animals (10.35 ± 3.23 vs. 18.00 ± 2.13%, *p* = 0.005; Fig. 3C). Since these values may also be influenced by potentially different AAV infectivity rates among granule cells (DAPI) or varying efficiency of the reporter induction (mCherry), we sought to assess whether these reactivation rates in GFP and 4D mice were significantly different from those expected from chance alone.

**Figure 3:**
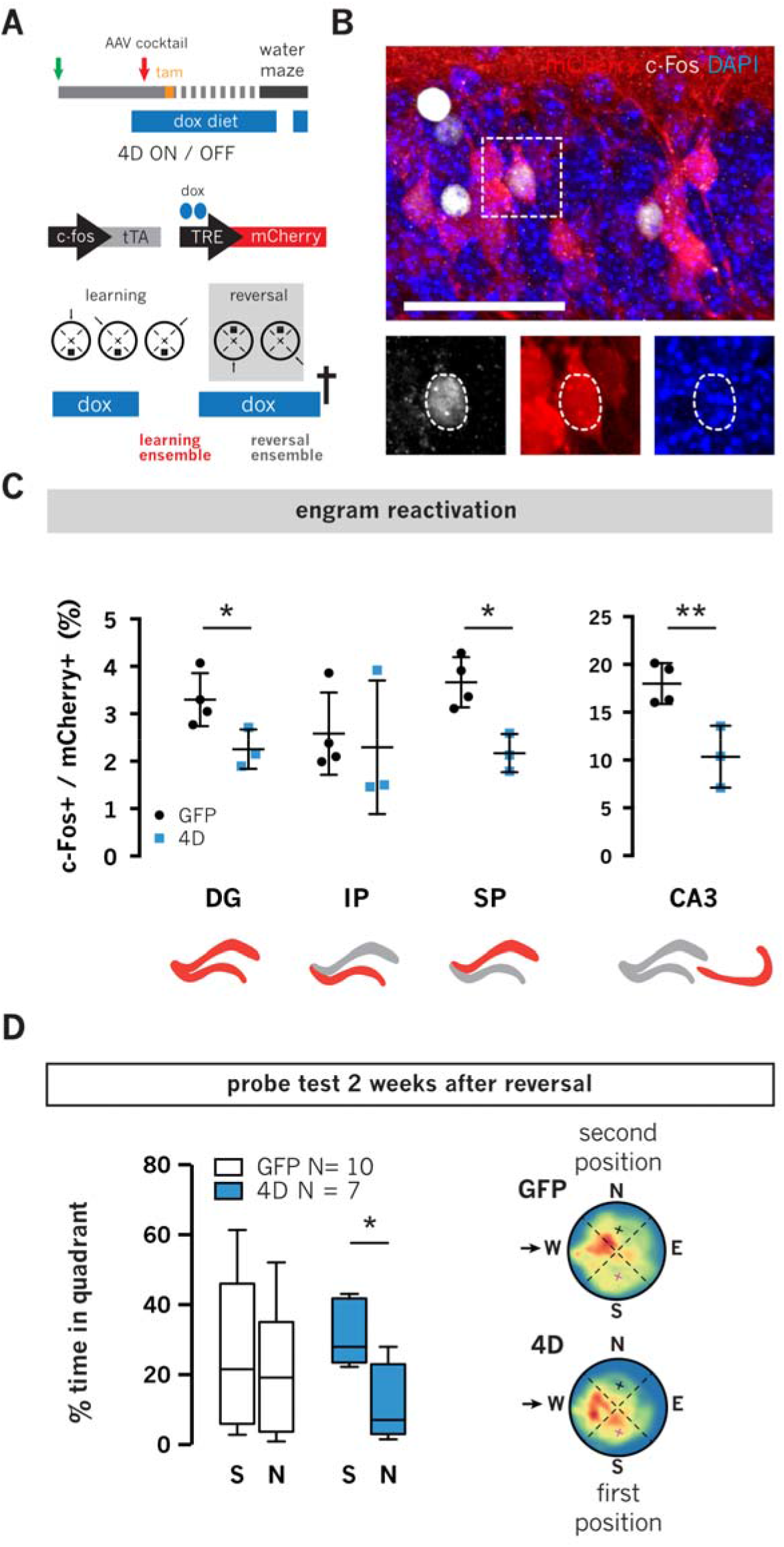
Increased neurogenesis facilitates the separation of memory indexes in the SP-CA3 circuit. **A-C)** Experimental layout (A), fluorescence pictures of engram-labelling (B) and quantifications (C) combining the use of AAV9 viruses encoding for *c-Fos*::tTa and *TRE*::mCherry with dox treatment for the labelling of remote (mCherry+) and recent (c-Fos+) memory representations. Note the reduction in reactivation rate specifically within the SP and CA3 areas (C). These data are also depicted relative to chance reactivation in Fig. S3B. **D)** Box-whisker plots (left) and heatmaps (right) displaying quadrant preference of mice tested 2 weeks after treatment shown in A. Note the increased preference of mice with increased neurogenesis for the first position of the platform. Scale bar = 100 μm. Data represent mean ± SD as the proportion of reactivated cells within the mCherry+ population in different DG-CA3 compartments (C) or box-whisker plots (D) of 4D (blue) and controls (black). N = 3-4 (C) and 7-10 (D); n > 400 (IP/SP) and > 100 (CA3). Unpaired (C) or paired (D) Student’s t-test * *p*<0.05; ** *p*<0.01.

To this aim, we calculated the probability of chance co-labelling (*P*_mCherry+_ * *P*_c-Fos+_ relative to DAPI+) in individual mice and found that while total reactivation in the SP blade was significantly above chance in control (0.30 ± 0.06 vs. 0.17 ± 0.09%, *p* = 0.006) it was less so and not significantly in 4D mice (0.20 ± 0.05 vs. 0.12 ± 0.01%, *p* = 0.085; Fig. S3B). This effect was also mirrored in the CA3 with chance levels of reactivation in control (4.06 ± 0.27 vs. 5.66 ± 2.69%, *p* = 0.28) and, remarkably, even below chance in 4D mice (1.29 ± 0.73 vs. 1.63 ± 0.69, *p* = 0.040; Fig. S3B). In turn, this is consistent with the notion that neurogenesis promotes the separation of memory indexes by increasing the sparsity of neural representations and thereby reduce the mere statistical probability that any given neuron is being re-activated twice in consecutive experiences (McAvoy et al., 2015). Hence, we expected that a strong positive correlation would emerge when comparing c-Fos levels and reactivation rates within the SP blade.

However, no obvious correlation could be found in neither control, 4D mice or pulling them together (*r* = 0.45, *p* = 0.30; Fig. S3C) and even the single mouse among controls showing the lowest level of c-Fos after reversal still displayed the highest level of SP reactivation (ca. 25% lower and 16% higher than the group mean, respectively; Fig. S3C, red dot). Intrigued by this unexpected lack in correlation within the SP, we next calculated the density of c-Fos+ (*P*_c-Fos_) cells that would be required to reduce total reactivation of GFP mice to their chance levels, as it was the case for 4D-treated animals, and resulting in a mathematically extrapolated reduction by 42.3 ± 19.5%, (i.e. from 1.71 ± 0.34 to 0.98 ± 0.09%). Surprisingly, this value was greater (*p* = 0.043) than the measured 13.9 ± 5.06% reduction in the density of c-Fos+ cells in the SP of 4D mice relative to controls (i.e. from 1.71 ± 0.34 to 1.47 ± 0.09%). In other words, the extrapolated reduction in c-Fos necessary for control mice to reach chance levels was significantly greater than the reduction observed in 4D mice. In turn, this suggested that neurogenesis can decrease reactivation rate in the SP beyond a mere reduction in the statistical probability of co-activation due to sparsity alone.

Finally, since neurogenesis has also been proposed to resolve interference through the forgetting of conflicting old memories (Epp et al., 2016), we investigated if the increased flexibility of 4D mice was also accompanied by attenuation of the memory of the first, learnt position of the platform by a probe trial 2 weeks after reversal. While GFP mice showed no preference for neither the first nor the second platform position (time in quadrants: 26.26 ± 21.73 vs. 20.13 ± 17.41%, respectively, *p* = 0.62), mice with increased neurogenesis preserved a clear preference for the first platform position (31.64 ± 8.94 vs. 12.18 ± 10.28%, *p* = 0.031; Fig. 3D). This indicated that the reduction of interference in animals with increased neurogenesis was not associated with forgetting of the first experience.

Altogether, our data showed that increasing neurogenesis allowed mice not only to learn more precisely but also to better detect novelty while at the same time retaining past memories. Our study further suggested that these effects were achieved through the recruitment of larger ensembles in the IP and the better separation of memory indexes in the SP, respectively. The distinct anatomical allocation and functional role of adult-born neurons within the blades of the DG may in turn improve memory precision and indexing that together combine to promote cognitive flexibility (Fig. 4).

**Figure 4:**
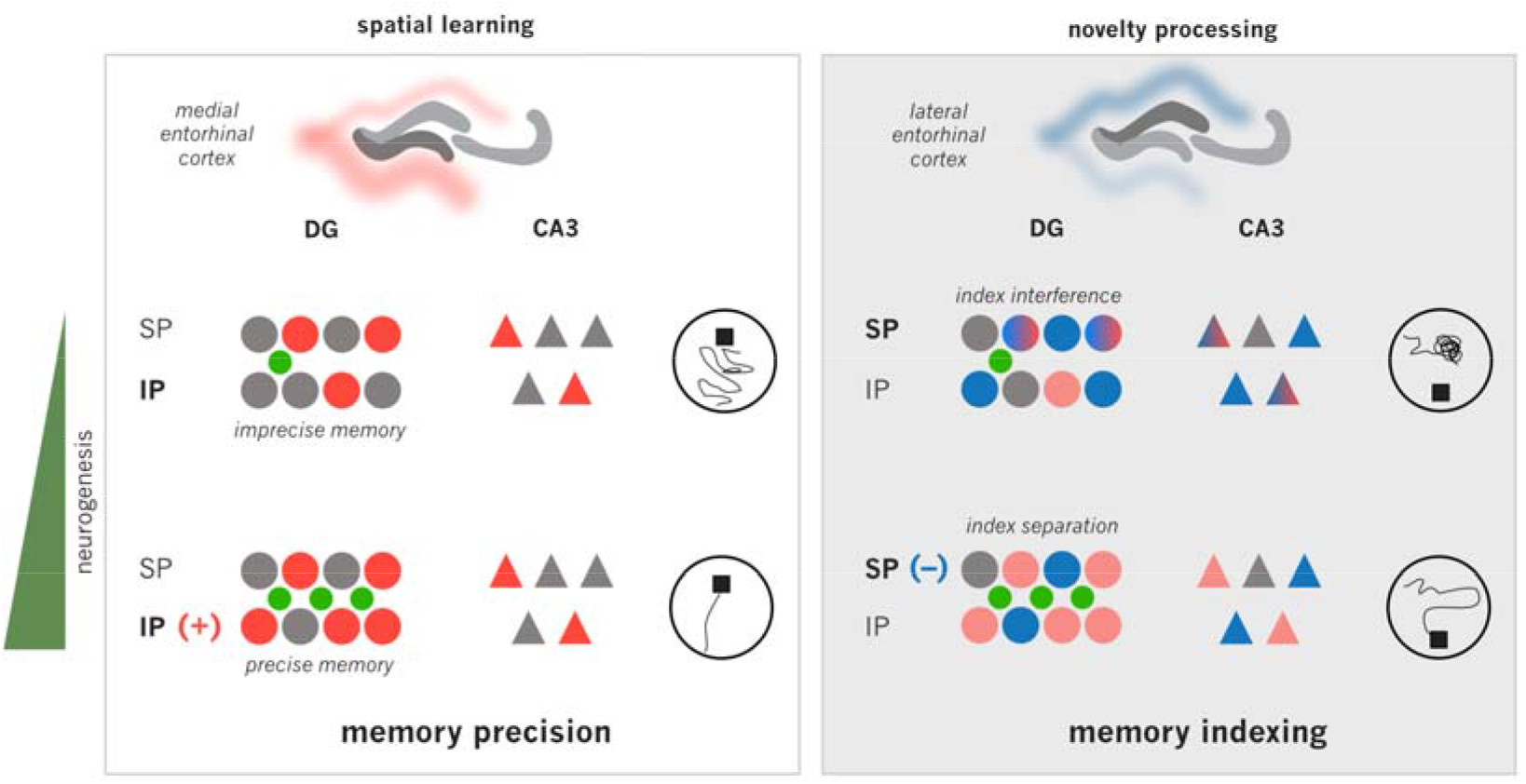
Model of input integration and DG blades regulation by adult-born neurons in cognitive flexibility. Together with a number of previous studies (see text), our study proposes a new model thereby input from the medial vs. lateral entorhinal cortex and carrying spatial vs. novelty information is differentially modulated by adult-born neurons of the IP vs. SP blades of the DG, which in turn is translated into gains in memory precision vs. indexing, respectively.

## DISCUSSION

Here we found that a cell intrinsic, genetic expansion of NSC increasing hippocampal neurogenesis differentially modulated IP-SP-CA3 activity during learning and improved cognitive flexibility. Importantly, the use of a previously characterized 4D model (Berdugo-Vega et al., 2020; Bragado Alonso et al., 2019) provided us with an unparalleled level of specificity for gain-of-function studies by the generation of a single cohort of age-matched neurons. This led us to address specific roles of 4 week old neurons in promoting memory precision and indexing through their differential regulation of the activity of the IP and SP blades of the DG. By looking at different phases of learning versus the introduction of novelty during navigation we further identified the activity patterns in the IP and SP that correlated with improvements in memory precision and indexing.

Specifically, while neurogenesis did not seem to have major effects in the progression from non-hippocampal to spatial navigation, it however promoted more spatially precise trajectories once hippocampal navigation was acquired. This behavioral gain was associated with an increased activity of the IP blade that did not directly involve the newborn neurons themselves. Knowing that mature granule cells are more spatially tuned than newborn neurons (Danielson et al., 2016), our study suggests that the effect of neurogenesis in recruiting a larger neuronal ensemble within the IP provides a better, more detailed spatial representation and improved memory precision (Fig. 4, left).

Moreover, and consistent with previous reports (Burghardt et al., 2012; Garthe et al., 2009, 2016), upon the introduction of novelty animals with increased neurogenesis displayed higher levels of flexibility. However, the nature of this flexibility and its significance at the level of memory representations remained elusive. In fact, to the best of our knowledge only two previous studies investigated the effects of neurogenesis in the separation of memory engrams during contextual discrimination (Denny et al., 2014; McAvoy et al., 2016). Yet, without discriminating local effects within the IP vs. SP blades and without separating behavioral effects in terms of spatial precision vs. novelty detection. We showed that neurogenesis-dependent sparsity in the SP blade after the introduction of novelty reduced the overlap in SP-CA3 memory representations, which is fundamental to achieve a better resolution of memory interference. Supporting the notion that newborn neurons act as novelty detectors (Denny et al., 2012), neurogenesis may facilitate the allocation of novel experiences to new or more distinct neuronal ensembles in the DG, CA3 or both (Denny et al., 2014; McAvoy et al., 2015; Niibori et al., 2012). Since the reduction in SP-CA3 activity in our study was not mediated by feed-forward inhibition as indicated by the similar levels of active, c-Fos+, parvalbumin+ interneurons in the CA3 (not shown), we infer that the reduced activity and reactivation rate of the CA3 is likely a direct result of changes in the SP that together dictate the recruitment of non-overlapping SP-CA3 memory ensembles. Reminiscent of the index theory (Teyler and DiScenna, 1986) and consistent with the proposed role of adult neurogenesis in increasing the capacity of memory indexing (Miller and Sahay, 2019), our study confines this role to 4 week old, newborn neurons within the SP (Fig. 4, right).

Finally, improvements in memory precision and indexing after increased neurogenesis did not occur at the cost of forgetting. This is fundamental and an intrinsic component of cognitive flexibility as the ability to acquire new memories without forgetting past ones. Intriguingly, control animals reacted to the introduction of novelty by promoting DG-CA3 activity, likely increasing the chance of interference between newly and previously formed memories. In contrast, a reduction in DG-CA3 activity occurring in conditions of increased neurogenesis may reduce such interference and as a result help forming new, while also preserving past, memories. It was surprising to observe that such effect of neurogenesis appeared to be even more potent than what deduced from the reduced statistical probability due to sparsity alone.

Adult neurogenesis has long been proposed to promote flexibility in learning but the underlying cognitive processes underlying such flexibility remained elusive. Our study dissects the anatomical and functional contribution of newborn neurons in promoting the formation of more precise memories in the IP and their better indexing by the SP blade, and provides insight into how plasticity modulates brain function to cope with an ever-changing environment.

## ACKNOWLEDGEMENTS AND AUTHORS CONTRIBUTION

We thank Simon Hertlein for technical assistance and Dr. Tomás Ryan for support with engram labelling experiments. This work was supported by the CRTD, TU Dresden, DFG CA893 17-1, a DIGS-BB fellowship awarded to GBV and EU-H2020 Marie Skłodowska-Curie grant (813851) to CCL. GBV and FC conceived the project, analysed the data and wrote the manuscript. AG and GK established analyses of navigational strategies and supported their interpretation. GBV performed all experiments supported by CCL. All authors contributed and approved the final manuscript. Animal experiments were approved by local authorities (TVV 39/2015 and TVV 56/2018). The authors declare no conflict of interest in this study.

## MATERIALS AND METHODS

### Viral preparations

Lentiviruses were produced by polyethyleneimine co-transfection of 293T cells with the respective transfer vectors encoding for GFP or 4D, HIV-1 gag/pol and VSV-G as previously described (Artegiani et al., 2011, 2012; Berdugo-Vega et al., 2020). 4D viruses encoded for the three transgenes (GFPnls/Cdk4/cyclinD1) linked by 2A peptides and with LoxP sites allowing the recombination of the 4D cassette together with the nls of GFP. Similar, floxed-nls-GFP vectors were used for control viral particles. One day after transfection, cells were switched to serum free medium and 1 day later the filtered supernatants were centrifuged at 25,500 rpmMice were kept in standard for 4 h. The viral particles were suspended in 40 μl of PBS per 10 cm petri dish and further concentrated using centrifugal filters (Amicon) yielding ca. 40 μl of virus suspension per construct with a titer of 10^8^-10^9^ IU/ml as assessed on HEK cells. A video of this protocol is available (Artegiani et al., 2012). AAV9 viruses for the reactivation experiment were produced by CaPO_4_ co-transfection of AAV293 cells (Stratagene, La Jolla) with the pAAV-cFos::tTA or pAAV-TRE::mCherry (gifts from William Widsen, Addgene plasmids #66794 and #66795), pHelper and pCR9. Cells were switched to fresh medium 8 h later and collected 2 days thereafter for freeze/thaw lysis. AAV particles were purified from the cell lysate by iodixanol gradient centrifugation (63,000 rpm for 2 h) and then washed in PBS and concentrated (Amicon filters) in successive centrifugation steps (4,000 g for 1 min). Viral titer was assessed by qPCR (10^10^ genome copies/ml).

### Animal treatments

Mice were kept in standard cages with a 12 h light cycle and water and food ad libitum. In all experiments, female *nestin*::CreRTt2 (Imayoshi et al., 2006) mice in a C57BL/6j genetic background were used. Briefly, isofluorane-anaesthetized mice were stereotaxically injected with 1 μl per hemisphere of HIV viral suspension in the DG hilus as previously reported (Artegiani et al., 2012) using a nanoliter-2000 injector (World Precision Instruments) and a stereotaxic frame Model 900 (Kopf Instruments) at ±1.6 mm mediolateral, −1.9 anteriorposterior, and −1.9 mm dorsoventral from bregma with a constant flow of 200 nl/min. For reactivation experiments, animals were switched to doxycycline diet (dox, 40 ppm, ENVIGO) 4 days before viral injections. AAV9 mixtures were obtained by dilution of c-Fos::tTA and TRE::mCherry viruses (1:20) and injected in the DG hilus. Dox diet was replaced by regular diet 24 h before the last day of learning and reintroduced after the last trial. Recombination of the 4D cassette and nls of GFP was achieved by oral administration of tamoxifen (Sigma) dissolved in corn oil (1:10) at 500 mg/kg body weight once a day for 3 non-consecutive days. BrdU (Sigma) diluted in PBS was intraperitoneally injected at 50 mg/kg concentration. Animals were either subjected to the water maze and/or anesthetized with pentobarbital and transcardially perfused with saline followed by 4% paraformaldehyde (PFA) fixation in phosphate buffer. Perfusions for c-Fos quantifications were performed 90 min after the last trial of navigation.

### Cellular quantifications

Brains were post-fixed overnight in 4% PFA at 4 °C and cut in coronal 40 μm thick vibratome sections serially collected along the rostro-caudal axis of the hippocampus and stored at −20 °C in cryoprotectant solution (25% ethylene glycol and 25% glycerol in PBS). Immunohistochemistry was performed stereologically in 1 every 6 sections (i.e. 10-12 sections analysed per brain) after blocking and permeabilization with 10% donkey serum in 0,3% Triton X-100 in PBS for 1,5 h at room temperature. Primary and secondary antibodies (Table S1) were incubated in 3% donkey serum in 0,3% Triton X-100 in PBS overnight at 4 °C. For BrdU detection sections were exposed to HCl 2 M for 25 min at 37 °C. DAPI was used to counterstain nuclei. Pictures were acquired using an automated Zeiss ApoTome or confocal (LSM 780) microscopes (Carl Zeiss) and maximal intensity projections of three optical sections (10 μm thick in total) taken and quantified using Photoshop CS5 (Adobe) and Affinity Photo. In all cases, GFP immunohistochemistry was performed to assess infectivity, which was confirmed in the vast majority (>90%) of the cases. Quantifications were performed on one series of DG sections per animal considering the subgranular zone (for NSC) or the whole thickness of the granular cell layer (for immature and mature neurons) of both the dorsal and ventral hippocampus. Neuronal activity was quantified in the IP and SP blades of the DG in the dorsal hippocampus (from −1.3 to −2.3 from Bregma) and reported as total number of c-Fos+ cells or density relative to DAPI (*P*_c-Fos_, *P*_mCherry_), extrapolated for the whole anatomical region being considered. Reactivation rate and total reactivation refer to the proportion of c-Fos+mCherry+ within the mCherry+ labelled cells and over the estimated number of DAPI, respectively, whereas chance reactivation was calculated as the intersection of individual probabilities for a given DAPI cell to be c-Fos+ and mCherry+ (% *P*_c-Fos_ * *P*_mcherry_). Areas were measured using Fiji 1.45b (ImageJ) and morphometric analyses performed on confocal reconstructions through the entire section to include most processes that were later traced and visualized using Fiji (ImageJ).

### Assessment of navigation

Behavioural experiments were performed in a Morris Water Maze located in a noise-isolated room with strong illumination (300 lux) and consisting of a learning (3 days) and reversal (2 days) phases, 6 trials per day as previously described (Garthe et al., 2009). Briefly, after acclimatization to the testing room (15 min) sessions were tracked using the Ethovision system (Noldus). At the end of each trial mice were guided to the platform and then placed in an individual drying cage. In the reversal phase the platform was relocated at 180 degrees. Probe tests consisted of a single trial for 1 min without platform. Navigational strategies were assessed by unbiased and automatic analysis of the swimming trajectories (Rtrack, Overall et al., 2020) and assigning each trial one predominant strategy among non-hippocampal, hippocampal (directed, focal and direct) or perseverance. Hippocampal strategies were further divided into precise and imprecise as a function of pathlengths (within or above a 1.5 threshold from optimal performance, respectively) and the initial trajectory error (the remaining distance left after a mouse has completed a path equal to the minimal trajectory) calculated using the Rtrack software. Quadrant occupancy was calculated using Ethovision.

### Statistics

Quantification of cell types and morphometric analyses were performed on independent biological replicates (see respective figure legends for N = mice and n = cells/dendritic segments/spines/trials counted per mouse) and depicted as means ± SDs or box-whisker plots with significance being assessed by two-tailed, unpaired (different animals) or paired (IP vs. SP and total vs. chance reactivation) Student’s t-test or Pearson’s correlation. Behavioural analyses were performed on 15-35 mice per group (as indicated in figure legends) and data depicted as area charts (navigational strategies) or box-whisker plots (pathlengths and probe tests). Statistical significance was accordingly calculated by Wald test of odds ratios assessed by logistic regression (navigational strategies), Mann-Whitney test (pathlengths of hippocampal strategies) or two-tailed paired (quadrant occupancy) or unpaired (performance) Student’s t-test.

---

**Supplementary Table 1.** List of primary antibodies

| antigen | dilution | manufacturer | catalogue # |
| --- | --- | --- | --- |
| BrdU | 1:250 | Abcam | ab6326 |
| c-Fos | 1:1000 | Synaptic Systems | 226003 |
| Dcx | 1:100 | Santa Cruz Biotechnology | Sc-8066 |
| GFP | 1:500 | Thermo Fisher | A-11122 |
| RFP (mCherry) | 1:500 | Chromotek | 5F8 |
| NeuN (Fox3) | 1:500 | Abcam | Ab104225 |
| Pv | 1:5000 | Swant | pv235 |
| S100β | 1:1000 | Abcam | ab14688 |
| Sox2 | 1:500 | Santa Cruz Biotechnology | Sc-17320 |

**List of antibodies** Form left to right: antigen recognized, dilution used, provider and catalogue number of primary antibodies. In most cases, Alexa Fluor secondary antibodies (Jackson ImmunoResearch) diluted 1:1000 were used.

**Supplementary Figure 1.**
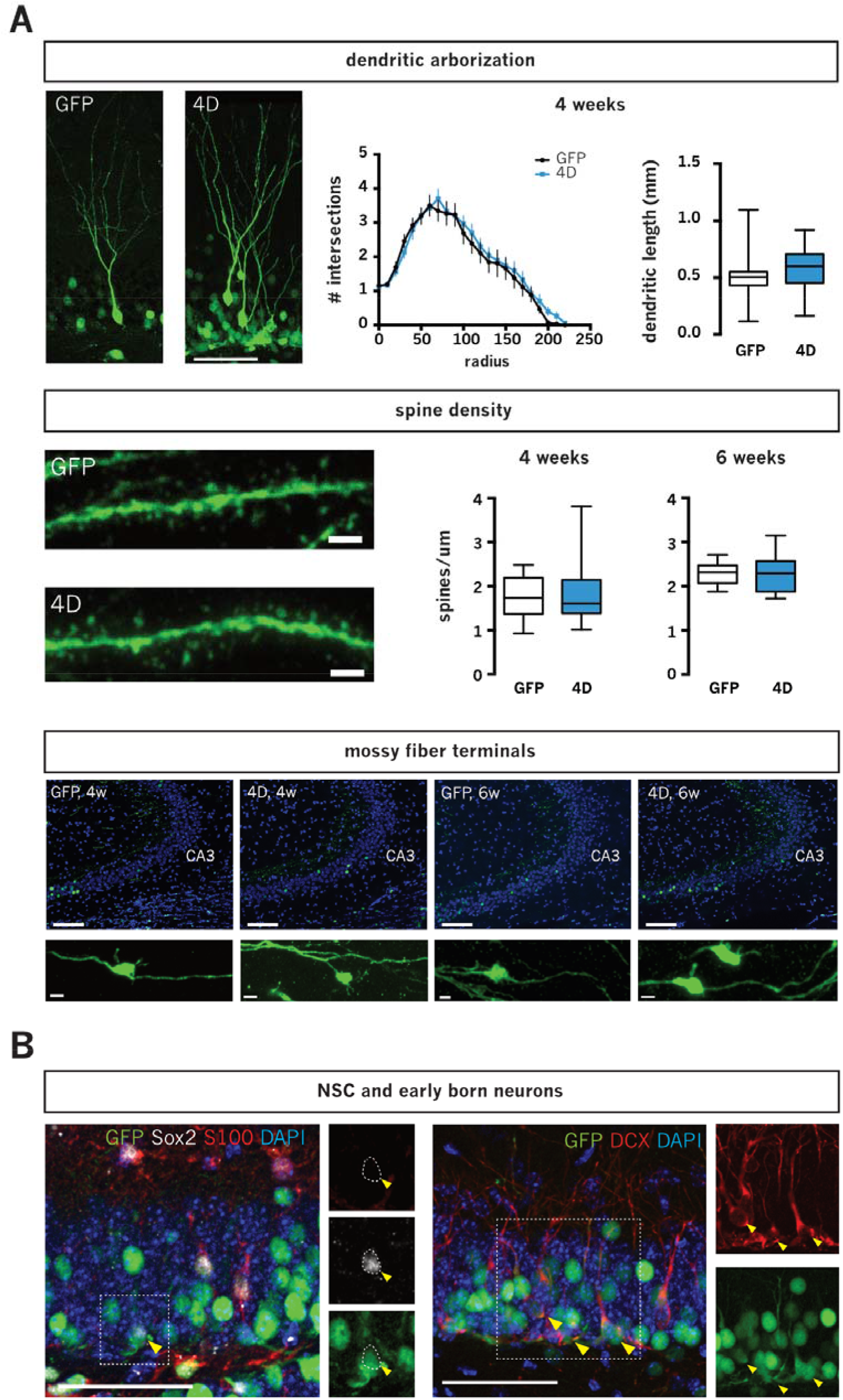
4D-generated neurons are morphologically normal. **A)** Fluorescence pictures and morphometric analysis of adult-born neurons of 4D-treated and control mice at different times after tamoxifen administration (as indicated) and identified by redistribution of GFP to the cytoplasm allowing the characterization of dendritic complexity and length (top), spine density (middle) and detection of axonal projection to the CA3 (bottom). **B)** Fluorescence pictures of cell types quantified in Fig. 1D and identified by markers of putative NSC (Sox2+/S100β−; left) and early born neurons (Dcx+; right) with examples (arrowheads). Scale bar = 100 μm (A; top and bottom, CA3; B) and 2 μm (A; middle and bottom, spines). Data represent mean ± SEM (Sholl profiles) and box-whisker plots (dendritic length and spines) for 4D (blue) and controls (white). N = 3; n > 26 (dendritic segments) and > 500 (spines from > 3 dendritic segments per animal).

**Supplementary Figure 2.**
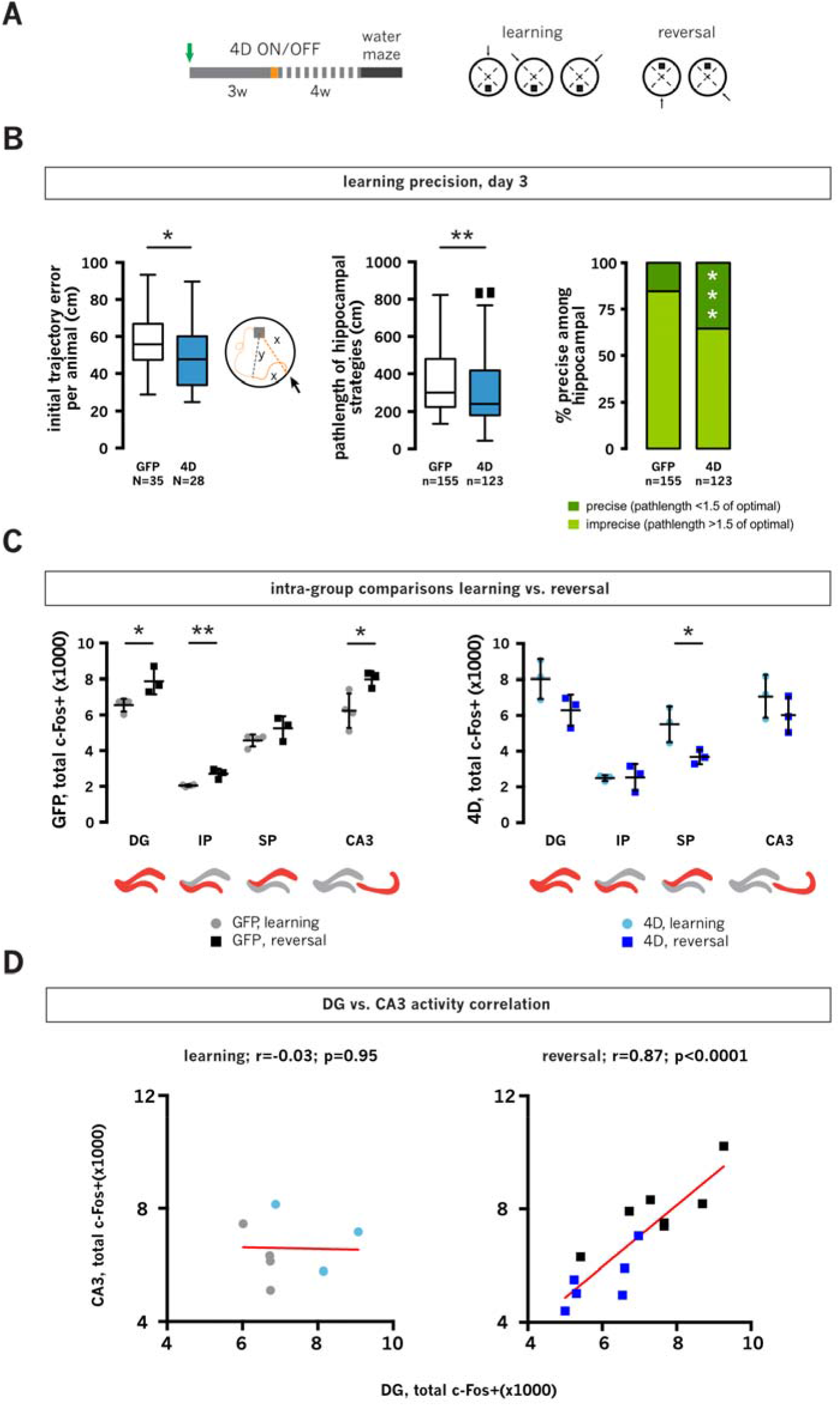
Increased neurogenesis promotes spatial memory precision and flexible learning. **A)** Experimental layout of GFP and 4D-treated animals subjected to the water maze and sacrificed after learning or reversal as in Fig. 2. **B)** Box-whiskers plots depicting initial trajectory error (left, depicted as “y” in scheme), pathlength among hippocampally-scored trajectories (middle) and bar graph representing the proportion of precise strategies (right, defined as relative to the optimal pathlength) of GFP (white) and 4D-treated (blue) animals on day 3 of learning. Note the improvements in all parameters in animals with increased neurogenesis. **C)** Intra-group analysis of c-Fos+ cells in DG compartments and CA3 (as indicated) calculated from data in Fig. 2D and grouping GFP and 4D animals after learning (light colors) and reversal (dark colors). Note the different response of GFP and 4D-treated mice to reversal. **D)** Pearson’s correlations of neural activity in the DG and CA3 after learning or reversal of the same animals shown in Fig. 2. N = 3-6 mice (C and D); n = 123-155 trials (B). Unpaired Student’s t-test (B, left and C), Mann-Whitney (B, middle) and Wald (B, right) tests * *p*<0.05; ** *p*<0.01.

**Supplementary Figure 3.**
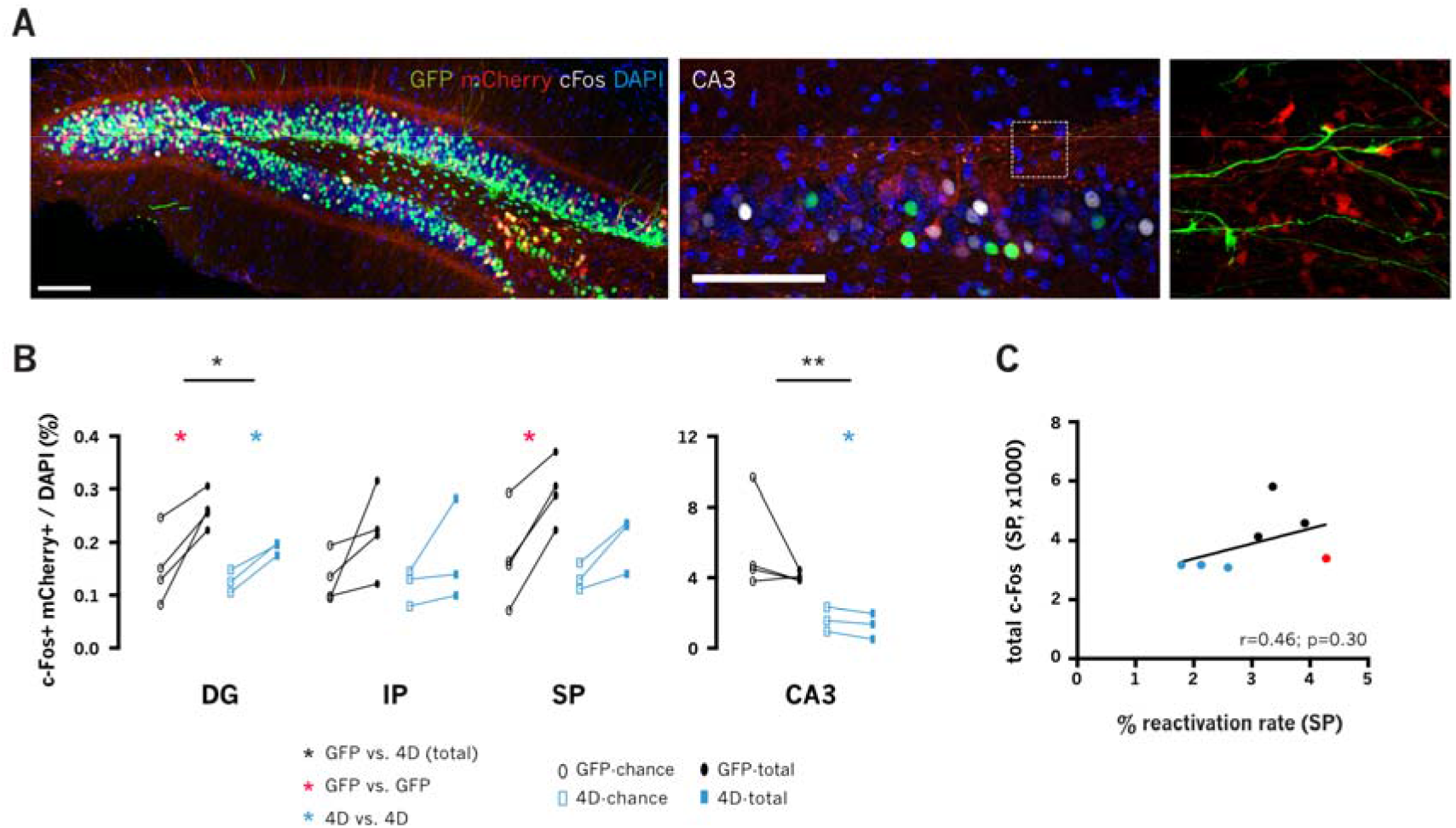
Cognitive flexibility is associated with the separation of SP-CA3 memory indexes. **A-B)** Fluorescence pictures with magnified insets (A) and quantifications of total and chance reactivation (B, as indicated) of animals shown in Fig. 3C. Data represent the relative changes in reactivation relative to chance per animal in DG compartments and CA3. **C)** Pearson’s correlation of SP activity with reactivation rate in the SP blade of the animals shown in Fig. 3C (N = 3-4, as indicated). Unpaired (inter-group) or paired (intra-group) Student’s t-test * *p*<0.05; ** *p*<0.01 as color coded.

## REFERENCES

Akers, K.G., Martinez-Canabal, A., Restivo, L., Yiu, A.P., Cristofaro, A.D., Hsiang, H.-L. (Liz), Wheeler, A.L., Guskjolen, A., Niibori, Y., Shoji, H., et al. (2014). Hippocampal Neurogenesis Regulates Forgetting During Adulthood and Infancy. Science 344, 598–602.

Anacker, C., and Hen, R. (2017). Adult hippocampal neurogenesis and cognitive flexibility — linking memory and mood. Nat. Rev. Neurosci. 18, 335–346.

Appleby, P.A., Kempermann, G., and Wiskott, L. (2011). The Role of Additive Neurogenesis and Synaptic Plasticity in a Hippocampal Memory Model with Grid-Cell Like Input. PLOS Comput. Biol. 7, e1001063.

Artegiani, B., Lindemann, D., and Calegari, F. (2011). Overexpression of cdk4 and cyclinD1 triggers greater expansion of neural stem cells in the adult mouse brain. J. Exp. Med. 208, 937–948.

Berdugo-Vega, G., Arias-Gil, G., López-Fernández, A., Artegiani, B., Wasielewska, J.M., Lee, C.-C., Lippert, M.T., Kempermann, G., Takagaki, K., and Calegari, F. (2020). Increasing neurogenesis refines hippocampal activity rejuvenating navigational learning strategies and contextual memory throughout life. Nat. Commun. 11, 1–12.

Bragado Alonso, S., Reinert, J.K., Marichal, N., Massalini, S., Berninger, B., Kuner, T., and Calegari, F. (2019). An increase in neural stem cells and olfactory bulb adult neurogenesis improves discrimination of highly similar odorants. EMBO J. 0, e98791.

Burghardt, N.S., Park, E.H., Hen, R., and Fenton, A.A. (2012). Adult-Born Hippocampal Neurons Promote Cognitive Flexibility in Mice. Hippocampus 22, 1795–1808.

Clelland, C.D., Choi, M., Romberg, C., Clemenson, G.D., Fragniere, A., Tyers, P., Jessberger, S., Saksida, L.M., Barker, R.A., Gage, F.H., et al. (2009). A Functional Role for Adult Hippocampal Neurogenesis in Spatial Pattern Separation. Science 325, 210–213.

Danielson, N.B., Kaifosh, P., Zaremba, J.D., Lovett-Barron, M., Tsai, J., Denny, C.A., Balough, E.M., Goldberg, A.R., Drew, L.J., Hen, R., et al. (2016). Distinct Contribution of Adult-Born Hippocampal Granule Cells to Context Encoding. Neuron 90, 101–112.

Denny, C.A., Burghardt, N.S., Schachter, D.M., Hen, R., and Drew, M.R. (2012). 4-to 6-week-old adult-born hippocampal neurons influence novelty-evoked exploration and contextual fear conditioning. Hippocampus 22, 1188–1201.

Denny, C.A., Kheirbek, M.A., Alba, E.L., Tanaka, K.F., Brachman, R.A., Laughman, K.B., Tomm, N.K., Turi, G.F., Losonczy, A., and Hen, R. (2014). Hippocampal Memory Traces Are Differentially Modulated by Experience, Time, and Adult Neurogenesis. Neuron 83, 189–201.

Dupret, D., Revest, J.-M., Koehl, M., Ichas, F., Giorgi, F.D., Costet, P., Abrous, D.N., and Piazza, P.V. (2008). Spatial Relational Memory Requires Hippocampal Adult Neurogenesis. PLOS ONE 3, e1959.

Epp, J.R., Silva Mera, R., Köhler, S., Josselyn, S.A., and Frankland, P.W. (2016). Neurogenesis-mediated forgetting minimizes proactive interference. Nat. Commun. 7, 10838.

Erwin, S.R., Sun, W., Copeland, M., Lindo, S., Spruston, N., and Cembrowski, M.S. (2020). A Sparse, Spatially Biased Subtype of Mature Granule Cell Dominates Recruitment in Hippocampal-Associated Behaviors. Cell Rep. 31, 107551.

Garthe, A., Behr, J., and Kempermann, G. (2009). Adult-Generated Hippocampal Neurons Allow the Flexible Use of Spatially Precise Learning Strategies. PLOS ONE 4, e5464.

Garthe, A., Roeder, I., and Kempermann, G. (2016). Mice in an enriched environment learn more flexibly because of adult hippocampal neurogenesis. Hippocampus 26, 261–271.

Hargreaves, E.L., Rao, G., Lee, I., and Knierim, J.J. (2005). Major Dissociation Between Medial and Lateral Entorhinal Input to Dorsal Hippocampus. Science 308, 1792–1794.

Kent, B.A., Beynon, A.L., Hornsby, A.K.E., Bekinschtein, P., Bussey, T.J., Davies, J.S., and Saksida, L.M. (2015). The orexigenic hormone acyl-ghrelin increases adult hippocampal neurogenesis and enhances pattern separation. Psychoneuroendocrinology 51, 431–439.

Kitamura, T., Saitoh, Y., Takashima, N., Murayama, A., Niibori, Y., Ageta, H., Sekiguchi, M., Sugiyama, H., and Inokuchi, K. (2009). Adult Neurogenesis Modulates the Hippocampus-Dependent Period of Associative Fear Memory. Cell 139, 814–827.

Knierim, J.J., Neunuebel, J.P., and Deshmukh, S.S. (2014). Functional correlates of the lateral and medial entorhinal cortex: objects, path integration and local–global reference frames. Philos. Trans. R. Soc. B Biol. Sci. 369.

Luna, V.M., Anacker, C., Burghardt, N.S., Khandaker, H., Andreu, V., Millette, A., Leary, P., Ravenelle, R., Jimenez, J.C., Mastrodonato, A., et al. (2019). Adult-born hippocampal neurons bidirectionally modulate entorhinal inputs into the dentate gyrus. Science 364, 578–583.

McAvoy, K., Besnard, A., and Sahay, A. (2015). Adult hippocampal neurogenesis and pattern separation in DG: a role for feedback inhibition in modulating sparseness to govern population-based coding. Front. Syst. Neurosci. 9.

McAvoy, K.M., Scobie, K.N., Berger, S., Russo, C., Guo, N., Decharatanachart, P., Vega-Ramirez, H., Miake-Lye, S., Whalen, M., Nelson, M., et al. (2016). Modulating Neuronal Competition Dynamics in the Dentate Gyrus to Rejuvenate Aging Memory Circuits. Neuron 91, 1356–1373.

Miller, S.M., and Sahay, A. (2019). Functions of adult-born neurons in hippocampal memory interference and indexing. Nat. Neurosci. 1–11.

Mishra, P., and Narayanan, R. (2020). Heterogeneities in intrinsic excitability and frequency-dependent response properties of granule cells across the blades of the rat dentate gyrus. J. Neurophysiol. 123, 755–772.

Niibori, Y., Yu, T.-S., Epp, J.R., Akers, K.G., Josselyn, S.A., and Frankland, P.W. (2012). Suppression of adult neurogenesis impairs population coding of similar contexts in hippocampal CA3 region. Nat. Commun. 3.

Overall, R.W., Zocher, S., Garthe, A., and Kempermann, G. (2020). Rtrack: a software package for reproducible automated water maze analysis. BioRxiv 2020.02.27.967372.

Piatti, V.C., Ewell, L.A., and Leutgeb, J.K. (2013). Neurogenesis in the dentate gyrus: carrying the message or dictating the tone. Front. Neurosci. 7.

Reijmers, L.G., Perkins, B.L., Matsuo, N., and Mayford, M. (2007). Localization of a Stable Neural Correlate of Associative Memory. Science 317, 1230–1233.

Rolls, E. (2013). The mechanisms for pattern completion and pattern separation in the hippocampus. Front. Syst. Neurosci. 7.

Sahay, A., Scobie, K.N., Hill, A.S., O’Carroll, C.M., Kheirbek, M.A., Burghardt, N.S., Fenton, A.A., Dranovsky, A., and Hen, R. (2011). Increasing adult hippocampal neurogenesis is sufficient to improve pattern separation. Nature 472, 466–470.

Teyler, T.J., and DiScenna, P. (1986). The hippocampal memory indexing theory. Behav. Neurosci. 100, 147–154.

Tonegawa, S., Pignatelli, M., Roy, D.S., and Ryan, T.J. (2015). Memory engram storage and retrieval. Curr. Opin. Neurobiol. 35, 101–109.

Wang, W., Pan, Y.-W., Zou, J., Li, T., Abel, G.M., Palmiter, R.D., Storm, D.R., and Xia, Z. (2014). Genetic Activation of ERK5 MAP Kinase Enhances Adult Neurogenesis and Extends Hippocampus-Dependent Long-Term Memory. J. Neurosci. 34, 2130–2147.

Wiskott, L., Rasch, M.J., and Kempermann, G. (2006). A functional hypothesis for adult hippocampal neurogenesis: Avoidance of catastrophic interference in the dentate gyrus. Hippocampus 16, 329–343.

